# Allele-specific control of replication timing and genome organization during development

**DOI:** 10.1101/221762

**Authors:** Juan Carlos Rivera-Mulia, Andrew Dimond, Daniel Vera, Claudia Trevilla-Garcia, Takayo Sasaki, Jared Zimmerman, Catherine Dupont, Joost Gribnau, Peter Fraser, David M. Gilbert

## Abstract

DNA replication occurs in a defined temporal order known as the replication-timing (RT) program. RT is regulated during development in discrete chromosomal units, coordinated with transcriptional activity and 3D genome organization. Here, we derived distinct cell types from F1 hybrid musculus X castaneus mouse crosses and exploited the high single nucleotide polymorphism (SNP) density to characterize allelic differences in RT (Repli-seq), genome organization (Hi-C and promoter-capture Hi-C), gene expression (nuclear RNA-seq) and chromatin accessibility (ATAC-seq). We also present *HARP*: a new computational tool for sorting SNPs in phased genomes to efficiently measure allele-specific genome-wide data. Analysis of 6 different hybrid mESC clones with different genomes (C57BL/6, 129/sv and CAST/Ei), parental configurations and gender revealed significant RT asynchrony between alleles across ~12 % of the autosomal genome linked to sub-species genomes but not to parental origin, growth conditions or gender. RT asynchrony in mESCs strongly correlated with changes in Hi-C compartments between alleles but not SNP density, gene expression, imprinting or chromatin accessibility. We then tracked mESC RT asynchronous regions during development by analyzing differentiated cell types including extraembryonic endoderm stem (XEN) cells, 4 male and female primary mouse embryonic fibroblasts (MEFs) and neural precursors (NPCs) differentiated *in vitro* from mESCs with opposite parental configurations. Surprisingly, we found that RT asynchrony and allelic discordance in Hi-C compartments seen in mESCs was largely lost in all differentiated cell types, coordinated with a more uniform Hi-C compartment arrangement, suggesting that genome organization of homologues converges to similar folding patterns during cell fate commitment.

## Introduction

Genome duplication in eukaryotes is regulated in coordination with large-scale chromosome organization and transcriptional activity, with discrete chromosome units (replication domains –RDs) replicating at specific times during S-phase (Hiratani et al. 2010; Rivera-Mulia and Gilbert 2016a; 2016b). Spatiotemporal regulation of RDs is critical for nuclear organization and function (Alver et al. 2014; Donley et al. 2015; Neelsen et al. 2013) and RT alterations are observed in disease (Ryba et al. 2012; Gerhardt et al. 2014b; 2014a; Dixon et al. 2017; Sasaki et al. 2017). Early and late RDs segregate to distinct nuclear compartments, with early replicating domains preferentially positioned at the nuclear interior while late RDs are located either at the periphery or close to the nucleolus (Jackson and Pombo 1998; Sadoni et al. 2004). Additionally, RDs correspond to the topologically-associating domains (TADs) measured by chromosome conformation capture techniques –Hi-C (Ryba et al. 2010; Moindrot et al. 2012; Yaffe et al. 2010; Pope et al. 2014; Rivera-Mulia and Gilbert 2016b). Moreover, RT is highly conserved in all eukaryotes (Ryba et al. 2010; Yue et al. 2014; Solovei et al. 2016) and changes dynamically during development in coordination with changes in nuclear positioning and transcriptional activity (Hiratani et al. 2010; Rivera-Mulia et al. 2015). Hence, RT constitutes a functional readout of genome organization and function.

Intriguingly, chromosome homologues replicate highly synchronously, with few exceptions that include imprinted genes (with the imprinted allele showing delayed replication) and mono-allelically expressed genes, which are also generally replicated earlier when active (Kitsberg et al. 1993; Simon et al. 1999; Mostoslavsky et al. 2001; Farago et al. 2012; Singh et al. 2003; Ensminger and Chess 2004; Dutta et al. 2009). X chromosome inactivation in female cells is the most impressive example of RT asynchrony linked to large-scale chromosome organization, with the inactive chromosome X (Xi) densely packed into the Barr body at the nuclear periphery and replicating later than the active chromosome X (Avner et al. 2001; Galupa and Heard 2015). Although most asynchronously replicating loci have been identified by cytogenetic analysis (Selig et al. 1992; Boggs and Chinault 1997), recent studies exploiting single nucleotide polymorphisms (SNPs) and deep sequencing of phased genomes allowed to measure allelic variation in RT genome-wide (Koren et al. 2014; Koren and McCarroll 2014; Mukhopadhyay et al. 2014; Bartholdy et al. 2015). These studies identified RT asynchrony associated with sequence variation and gene imprinting. Although differential efficiency in replication origin firing has been associated with allelic variation in RT (Bartholdy et al. 2015) it is not clear whether the RT asynchrony is linked to differences in 3D genome organization between alleles and if those differences are regulated during development. Here, we took advantage of the high SNP density between genomes of distinct mouse subspecies and measured allelic differences in RT, 3D genome organization, gene expression and chromatin accessibility. To do so we developed the algorithm *Haplotype-Assisted Read Parsing* (HARP), a new computational tool to efficiently sort reads into each genome based on the presence of SNPs. We identified RT asynchrony in ES cells that correlated with allelic differences in Hi-C compartments but not SNP density, gene expression or chromatin accessibility. Surprisingly, we found that RT asynchrony is lost during cell fate specification towards distinct cell types in coordination with a decrease in Hi-C compartment differences.

## Results

To evaluate allelic RT variation we exploited the high single nucleotide polymorphisms (SNPs) density between distinct mouse genomes. Mouse crosses of subspecies and strains were generated and ES cell lines were derived from hybrid F1 blastocysts (Figure 1A) as previously described (Dupont et al. 2016). We characterized a total of 6 hybrid mES cell lines harboring 3 different genomes (C57BL/6, 129/sv and CAST/Ei), opposite parental configurations and gender (Figure 1B). Genome-wide RT analysis (Figure 1C) was performed by NGS as described previously (Ryba et al. 2011; Marchal et al. 2017). Allele-specific genome-wide data was measured using our *HARP* algorithm (see Methods). Chromatin spatial organization (Hi-C) and long range enhancer-promoter interactions (PC-Hi-C), gene expression (nuclear RNAseq) and chromatin accessibility (ATACseq) data were also collected (see methods) to identify possible mechanisms driving the differences between alleles in RT and 3D genome organization (Figure 1D–F). An exemplary genomic region with allele-specific measurements of RT, Hi-C compartments and interactions, gene expression and chromatin accessibility is shown in Figure 1G.

**Figure 1.**
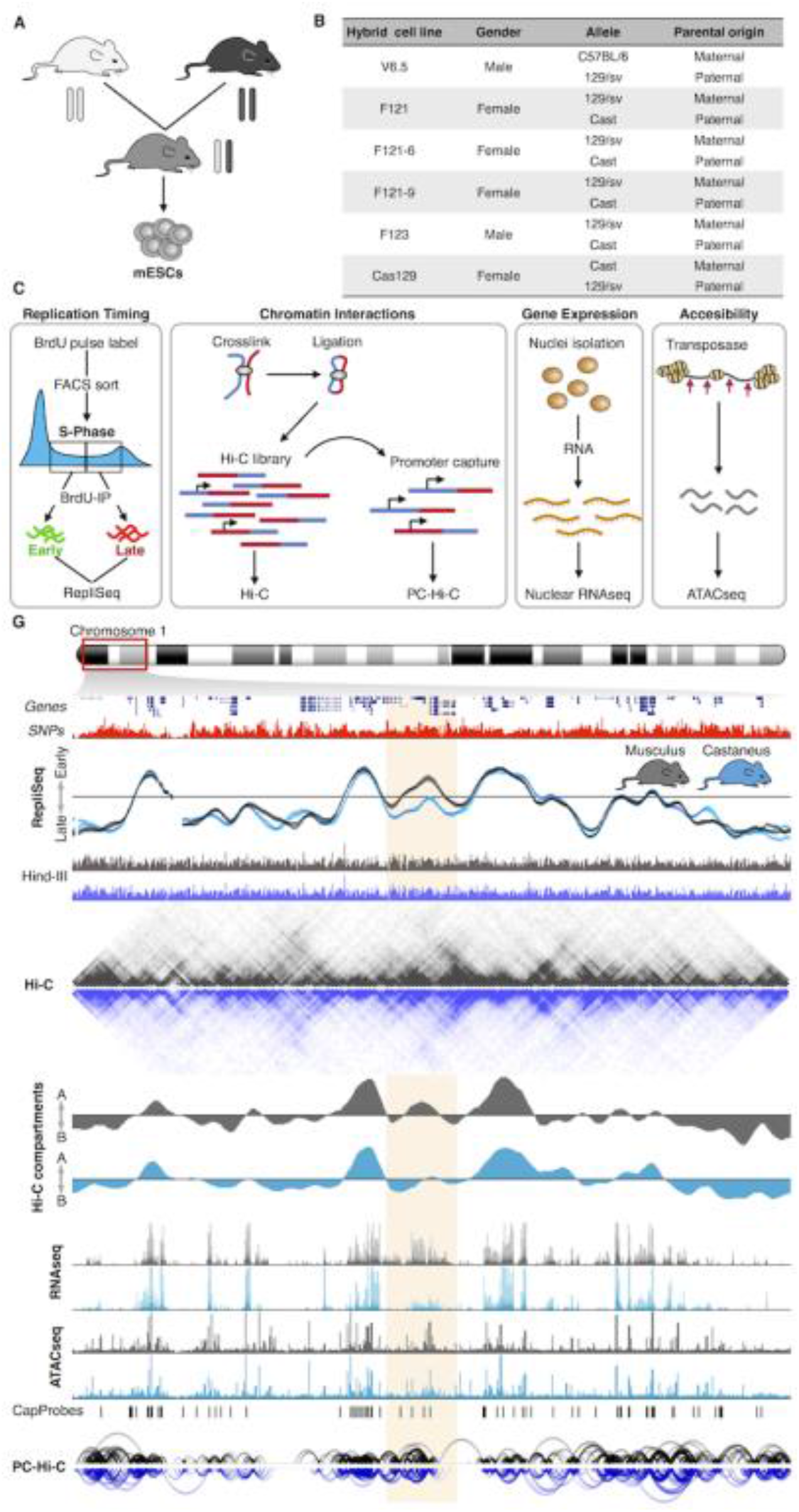
Genome-wide analysis of RT, enhancer-promoter interactions, gene expression and chromatin accessibility in hybrid mouse ESCs (click for full resolution). (A) Mouse ESC lines were derived from hybrid F1 blastocysts from crosses of distinct subspecies and strains. (B) Six distinct hybrid mES cell lines harboring 3 different genomes (C57BL/6, 129/sv and Cast), opposite parental configurations and different genders were analyzed. V6.5, F121 and F123 hybrid cell lines were generated previously (Monkhorst et al. 2008; Rideout et al. 2000). F121-6 and F121-9 are single cell sub-clones of F121 and Cas129 was generated in this study from a reciprocal cross between castaneus/musculus mice. C-F Genome-wide analysis of RT (C), Hi-C and promoter capture Hi-C [PC-Hi-C] (D), RNAseq (E) and chromatin accessibility measured by <u>A</u>ssay for <u>T</u>ransposase <u>A</u>ccessible <u>C</u>hromatin–ATAC-seq (F). (G) Representative genomic region on chromosome 1 showing RT profiles of musculus (129/sv) and castaneus (CAST/Ei) alleles. Two replicates of the F121-9 cell line (two RT profiles of each genome) show the consistency in RT asynchrony. Allele-specific Hi-C matrices and compartments A and B (eigenvectors), RNA-seq, ATAC-seq and enhancer-promoter interactions are shown. Allele-specific RT, RNAseq and ATACseq were determined based on the SNPs shown in red. Allele-specific Hi-C and PC-Hi-C interactions were obtained using only Hind-III fragments containing SNPs that distinguish each genome (Hind-III track). Capture probes for PC-Hi-C are shown above promoter-enhancer interactions. Hi-C data was obtained from (Giorgetti et al. 2016).

### RT asynchrony between alleles in hybrid mouse ESCs

To identify the degree of RT synchrony we divided the autosomal genome into 43,941 non-overlapping 50 kb windows. First, we determined the degree of similarity in RT between the two alleles. As expected we found the strongest genome-wide correlation in the ESC line derived from a mouse inbred cross (genomes C57BL/6 and 129/sv) and slight lower correlation values for ESCs derived from subspecies *M. musculus musculus* (129/sv genome) and *M. musculus castaneus* (CAST/Ei genome) crosses (Figure 2A). Interestingly, although the correlation values were very strong (>0.9) between the alleles from all hybrid mouse ES cell lines analyzed, they were segregated per sub-species in a genome-wide correlation matrix and the highest correlation values were observed between alleles with the same sub-species genome (either *M. musculus musculus* or *M. musculus castaneus)* rather than between alleles from the same cell line (Figure 2B). Moreover, alleles from the V6.5 cell line (derived from a cross of inbreeds from the same sub-species) clustered tightly together with the rest of the *M. musculus musculus* genomes (Figure 2B). Next, to identify the RT asynchrony between homologues, we analyzed the magnitude of differences in RT between alleles and between replicates. Considering an average *S*-phase time of 8hrs, we found that the largest differences between replicates of the same allele (same genome) were <80 minutes (Supplementary figure 1), which is consistent with the average technical noise commonly observed in RT measurements (approximately 10% of the dynamic range). Hence, we considered significant RT asynchrony between alleles to be any difference larger than 80 minutes (Figure 2C–D). In agreement with the correlation analysis, we found very few asynchronous regions in the V6.5 cell line (0.7%) with the *C57BL/6* and *129/sv* genomes (Figure 2C–E and Supplementary figure 2) but 12 % of the genome replicating asynchronously in cell lines derived from *M. musculus musculus* and *M. musculus castaneus* crosses (Figure 2C–E and Supplementary figure 3).

**Figure 2.**
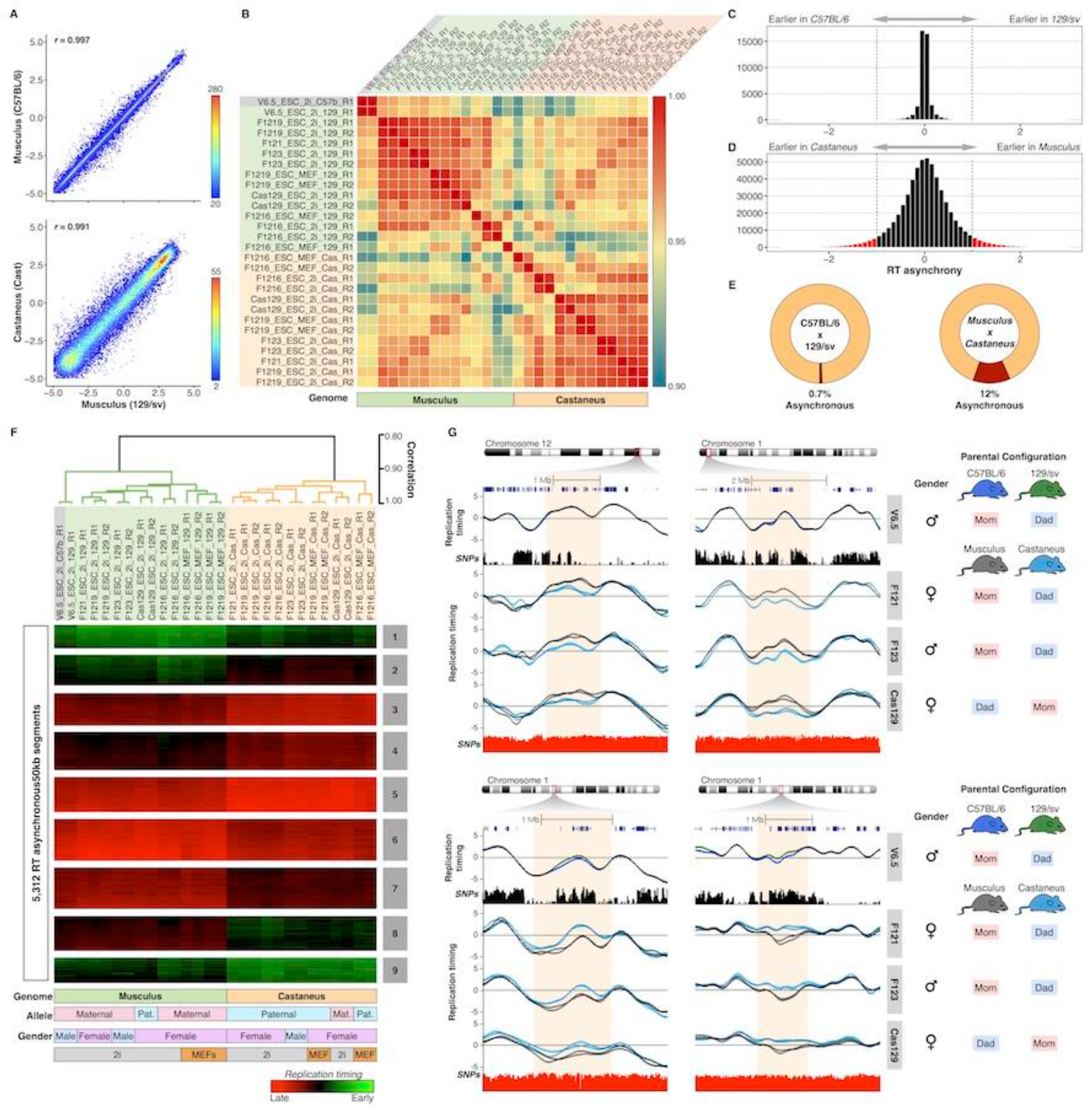
Replication timing asynchrony in hybrid mouse ESCs (<u>click for full resolution</u>). (A) Genome-wide correlations of RT between alleles for hybrid ESC lines V6.5 (top panel) and F1219 (bottom panel). RT values of 50kb windows were displayed as binned scatter plots with a density heatmap. (B) Correlation matrix of allele-specific RT separate samples by genome. Data from all cell lines (V6.5, F121, F1216, F1219, F123, Cas129) were analyzed and arranged based on their correlation values. C-D RT asynchrony in V6.5 and castaneus x musculus mES cell lines. Histogram of RT differences between C57BL/6 and 129/Sv (C) and between 129/sv and castaneus genomes (D) are shown. (E) Percentages of autosomal genome showing RT asynchrony in C57BL/6 x 129/sv and Musculus x Castaneus are shown. Asynchrony was defined as differences > 80 min in S-phase. (F) Unsupervised clustering analysis of RT asynchronous chromosomal segments identified specific RT signatures. Heat map shows the RT ratios [=log2(Early/Late)]. Branches of the dendrogram were constructed based on the correlation values (distance = correlation value –1) and a correlation threshold of 0.9 was used to define two main sample clusters. *k-means* clustering analysis of RT asynchronous regions defined RT signatures with specific patterns. Specific genomes, parental configuration, gender and growth conditions are shown at the bottom. (G) RT profiles of exemplary genomic regions showing RT asynchrony between alleles. Cell lines are labeled at the right in grey boxes, SNPs between C57B/6 vs. 129/sv and 129/sv vs. Cast are shown in black and red peak tracks respectively. C57B/6 alleles are shown as green lines, 129/sv alleles as grey lines and Castaneus as blue lines. Two replicates of each cell line are shown in each plot and the gender and parental configuration of each hybrid mESC line is shown.

Next, we extracted all genomic regions with replication asynchrony and classified them into RT signatures (Figure 2F–G) as previously described (Rivera-Mulia et al. 2015). As expected, the clustering analysis showed that alleles from the V6.5 cell line clustered together and with all other alleles from *Mus musculus musculus* genomes while *Mus musculus castaneus* alleles formed a separated cluster (Figure 2F). Unsupervised clustering confirmed that the allelic differences in RT are associated with their respective sub-species genomes but not to parental origin or gender (Figure 2F). In fact, only 0.10% of the total RT variation across all samples is linked to the gender of mouse ESCs (Supplementary figure 3) and no differences were observed linked to parental origin (Figure 2F). Additional analysis of mouse hybrid ESCs with opposite parental configuration (F121 and F123 compared to Cas129) confirmed that RT differences are associated to *musculus* and *castaneus* genomes but not parental origin (Figure 2G). Exemplary RT profiles show that the differences in RT between distinct alleles span over megabase regions and occur between sub-species genomes (Figure 2G).

### RT asynchrony in hybrid mouse ESCs occurs at developmentally regulated replication domains

We also analyzed whether the RT asynchronous regions identified in hybrid mouse ES cells coincide with replication domains that are regulated during development (i.e. domains that change between early and late compartments during distinct cell specification pathways). To do so, we identified developmentally regulated RT domains (RDs) across 30 mouse cell lines representing cell types from each of the three germ layers and measured the overlap with the hybrid mouse ESC asynchronously replicating regions. We found that 67.4 % of the RT asynchrony occurred at developmentally RT regulated genomic regions (Supplementary Figure 3). By using sequential Monte Carlo multiple testing (MCFDR) algorithm (Sandve et al. 2011; 2010), preserving the RDs lengths and positions and randomizing the positions of the RT asynchronous regions, we found a highly significant overlap as compared to what is expected by chance (p-value = 0.003984).

### Allelic differences in RT are maintained under different ES media conditions

RT asynchrony between *musculus* and *castaneus* genomes was maintained under different growth conditions: either when cultured on MEFs in serum + LIF or feeder-free in 2i media (Figure 2F and Supplementary Figure 4). In fact, we found that only 0.17% of the autosomal genome showed RT differences that correlated with growth conditions (Supplementary figure 3). Overall, our results suggest that allelic RT differences detected in ESCs are associated with the sub-species genomes but not with gender or paternal configuration and are stable in distinct growth conditions that maintain the naïve vs. ground states of pluripotency.

### RT asynchrony correlates with 3D genome organization but not sequence variation, gene expression or chromatin accessibility

Previous analysis in human cells suggested that sequence variation is linked to RT asynchrony between alleles (Koren et al. 2014; Mukhopadhyay et al. 2014; Bartholdy et al. 2015). Here, we analyzed the relationship between RT differences and SNP density to determine whether RT asynchrony was related to local SNP density. We found that RT asynchronous regions were not enriched for SNPs and no significant differences in SNP density were observed between RT constitutive regions and RT variable regions (Figure 3A).

**Figure 3.**
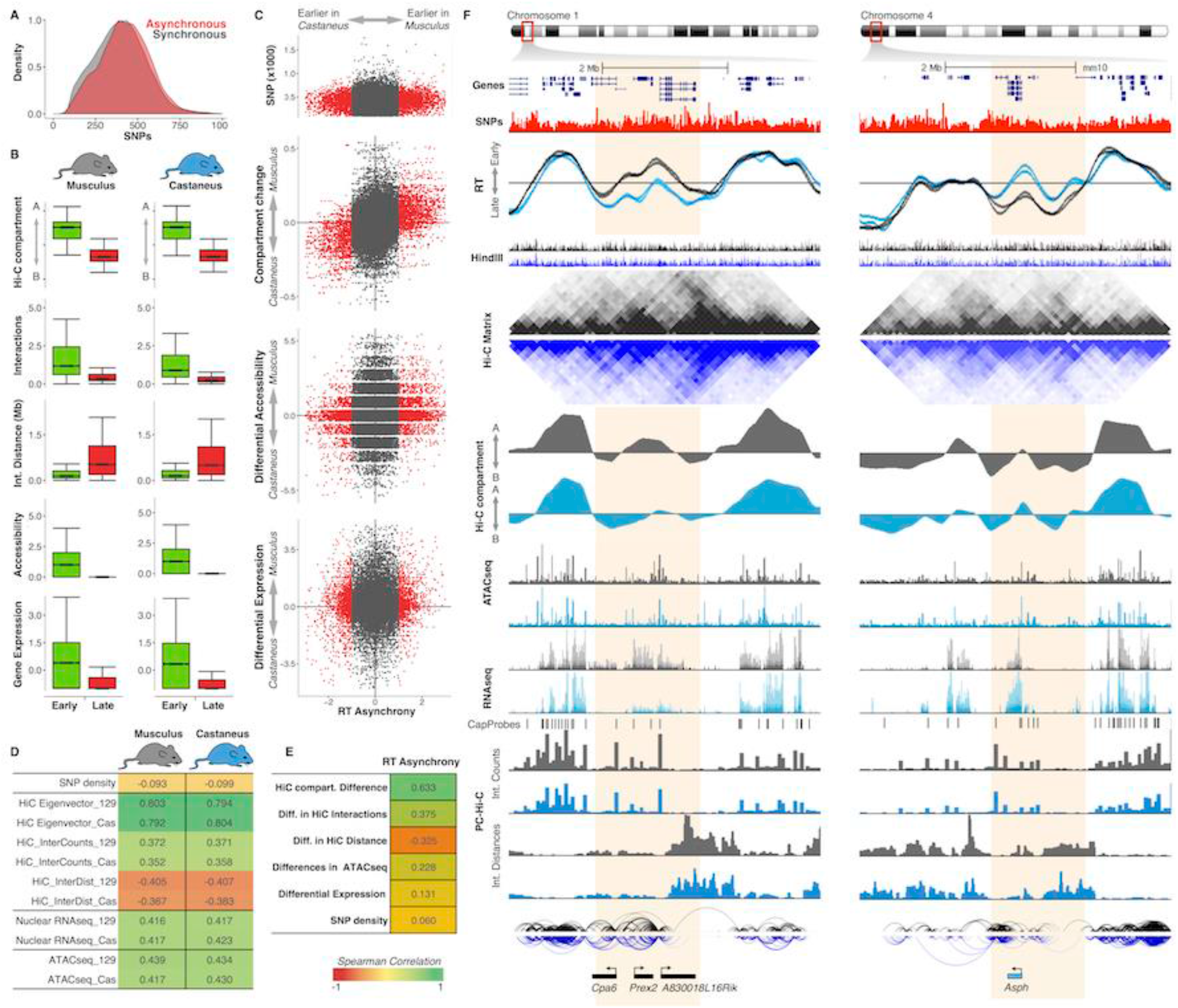
RT asynchrony correlates with genome organization (click for full resolution) (A) RT synchronous and asynchronous genomic regions contain similar SNP densities. (B) Genome organization, chromatin accessibility and gene expression of RT synchronous regions that replicate either early or late during S-phase. (C) Scatter plots of RT asynchrony vs. SNP density, changes in Hi-C compartments, differential accessibility and differential expression. Hi-C data was obtained from (Giorgetti et al. 2016). (D) Spearman correlation values of RT and distinct genomic features per genome. (E) Spearman correlation values of RT asynchrony and changes in Hi-C compartments, Hi-C interaction counts and distances, differences in chromatin accessibility and SNP density. (F) Two exemplary chromosome regions showing the RT asynchrony associated with changes in Hi-C compartments, differential expression and distinct promoter-enhancer interactions. Two replicates of each cell line are shown in each plot of RT profiles. Allele-specific RT, RNAseq and ATACseq were determined based on the SNPs shown in red. Allele-specific Hi-C and PC-Hi-C data was obtained using Hind-III fragments containing SNPs for each genome (Hind-III track). Differentially expressed genes measured by nuclear RNAseq are shown at the bottom color coded according to the allele showing the highest expression value (musculus = black, light blue = castaneus).

Next, we analyzed the global chromatin properties of the genomic regions in which the two alleles replicate synchronously in hybrid mouse ESCs, comparing chromosomal segments that replicate early vs. late during S-phase. Consistent with previous studies (Ryba et al. 2010; Pope et al. 2014; Rivera-Mulia and Gilbert 2016b), we found that early replicating regions are within Hi-C compartment A while late replicating regions are within Hi-C compartment B (Figure 3B). Moreover, synchronously early replicating allelic regions presented higher densities of long-range chromatin interactions with shorter distances in comparison with the synchronously late replicating allelic regions (Figure 3B). Similarly, we found that early replicating regions were more accessible and the genes located within them expressed at higher levels in comparison to late replicating regions (Figure 3B). Genome-wide correlation analysis of RT of each allele amongst the synchronously replicating portion of the genome with several genomic features confirmed that 3D chromatin organization (Hi-C) is the genomic property with the strongest correlation with RT (Figure 3D).

We next analyzed the correlation of the asynchronously replicating regions of the genome with all other genomic properties. Scatter plots and spearman correlation demonstrate that changes in 3D genome organization strongly correlated (correlation >0.6) with RT asynchrony (Figure 3C and 3E). Moreover, this analysis confirmed that SNP density was not linked to the RT differences between the chromosome homologues (Figure 3C and 3E). Surprisingly, the differences in chromatin accessibility and gene expression did not correlate with RT asynchrony (Figure 3C and 3E). Exemplary RT profiles of chromosomal regions with RT asynchrony confirm: 1) the close link between RT and 3D genome organization (Hi-C eigenvectors); 2) similar patterns of chromatin accessibility and gene expression between the two alleles and; 3) the differences in PC-Hi-C interaction counts and distances between early and late replicating compartments (Figure 3F).

### RT asynchrony in hybrid mouse ES cells is not associated with gene imprinting

Early observations of RT asynchrony linked differences in RT to gene imprinting (Reik and Walter 2001). Moreover, a recent genome-wide study of RT asynchrony in human adult erythroid cells suggested that allelic differences in RT are enriched in imprinted genes (Mukhopadhyay et al. 2014). Hence, we analyzed whether the RT asynchrony identified in hybrid mouse ES cells is associated with gene imprinting. We obtained RT values at the transcription start sites (TSS) of all RefSeq genes, identified the genes with RT asynchrony and measured their overlap with characterized imprinted genes. We found that only 1.5% of the RT asynchronous genes are imprinted genes (Supplementary Figure 5). Moreover, only a fraction (25.5%) of imprinted genes replicated asynchronously in hybrid mouse ESCs (Supplementary Figure 5). However, allelic differences in RT at those imprinted genes are not linked to gene imprinting as identical RT patterns were observed in cell lines with opposite parental configuration. Hence, RT asynchrony in hybrid mouse ESC lines is not due to parental imprinting (Supplementary Figure 5).

### Long-range interactions in RT asynchronous domains are restricted to the early replicating allele

RT and Hi-C eigenvectors correlated strongly and RT asynchrony was associated to differences in Hi-C compartments (Figure 3), suggesting that differences in chromatin interactions and 3D genome organization are linked to the allelic differences in RT. Hence, to test whether specific interactions are associated with the changes in RT between homologues we analyzed PC-Hi-C data, which reduces the complexity of Hi-C libraries and allows the identification of significant regulatory interactions (Mifsud et al. 2015; Schoenfelder et al. 2015a). We did not find significant differences when comparing the total number of interactions and average distances between the two homologues at the asynchronous replicating regions. However, we found that discrete long range enhancer-promoter interactions were restricted to the allele replicating earlier (Figure 3F and Supplementary Figure 6). In contrast, PC-Hi-C interactions at the allele replicating later in the asynchronous regions were restricted to short-range distances and the majority were within the replication domain (Figure 3F and Supplementary Figure 6). These observations suggest that long-range enhancer-promoter interactions connecting early replicating domains with RT asynchronous domains are restricted to the allele that replicates earlier.

### RT asynchrony is lost during cell fate commitment

To determine whether RT asynchrony is maintained or increased during loss of pluripotency, we generated allele-specific RT programs of extraembryonic endoderm stem (XEN) cells (Dupont et al. 2016) and four different primary mouse embryonic fibroblasts (MEFs) derived from *musculus* X *castaneus* F1 hybrid embryos, as well as for neural precursors (NPCs) differentiated *in vitro* from hybrid mouse ESC lines with opposite parental configuration (Figure 4A). Consistent with our previous findings demonstrating that RT is cell type specific (Hiratani et al. 2010; Rivera-Mulia et al. 2015), genome-wide correlation confirms different RT programs for the distinct hybrid cell types (Supplementary Figure 7). However, in contrast to ESCs, higher correlations between alleles, replicates and cell lines were observed for all differentiated hybrid cell types (MEFs, NPCs and XEN cells), suggesting fewer differences in RT between homologues (Supplementary Figure 7). Consistently, we found a dramatic decrease in RT asynchrony in all differentiated cell types: 6% in XEN cells, 4% in MEFs and 1% in NPCs of the autosomal genome. Moreover, very little overlap of RT asynchrony was observed between the distinct hybrid cell types, suggesting that allelic difference in RT are epigenetically regulated during development (Supplementary Figure 8); although these asynchronous regions were not associated with gene imprinting (Supplementary Figure 9). Furthermore, we tracked the RT of hybrid mouse ESC asynchronous regions in MEFs, NPCs and XEN cells and found that more than 85% of the regions with allelic RT differences in stem cells became synchronously replicated in all differentiated cells (Figure 4C). Exemplary RT profiles of hybrid mouse ESCs, XEN cells, MEFs and NPCs confirm that the regions of RT asynchrony observed in pluripotency are lost during cell fate commitment (Figure 4D).

**Figure 4.**
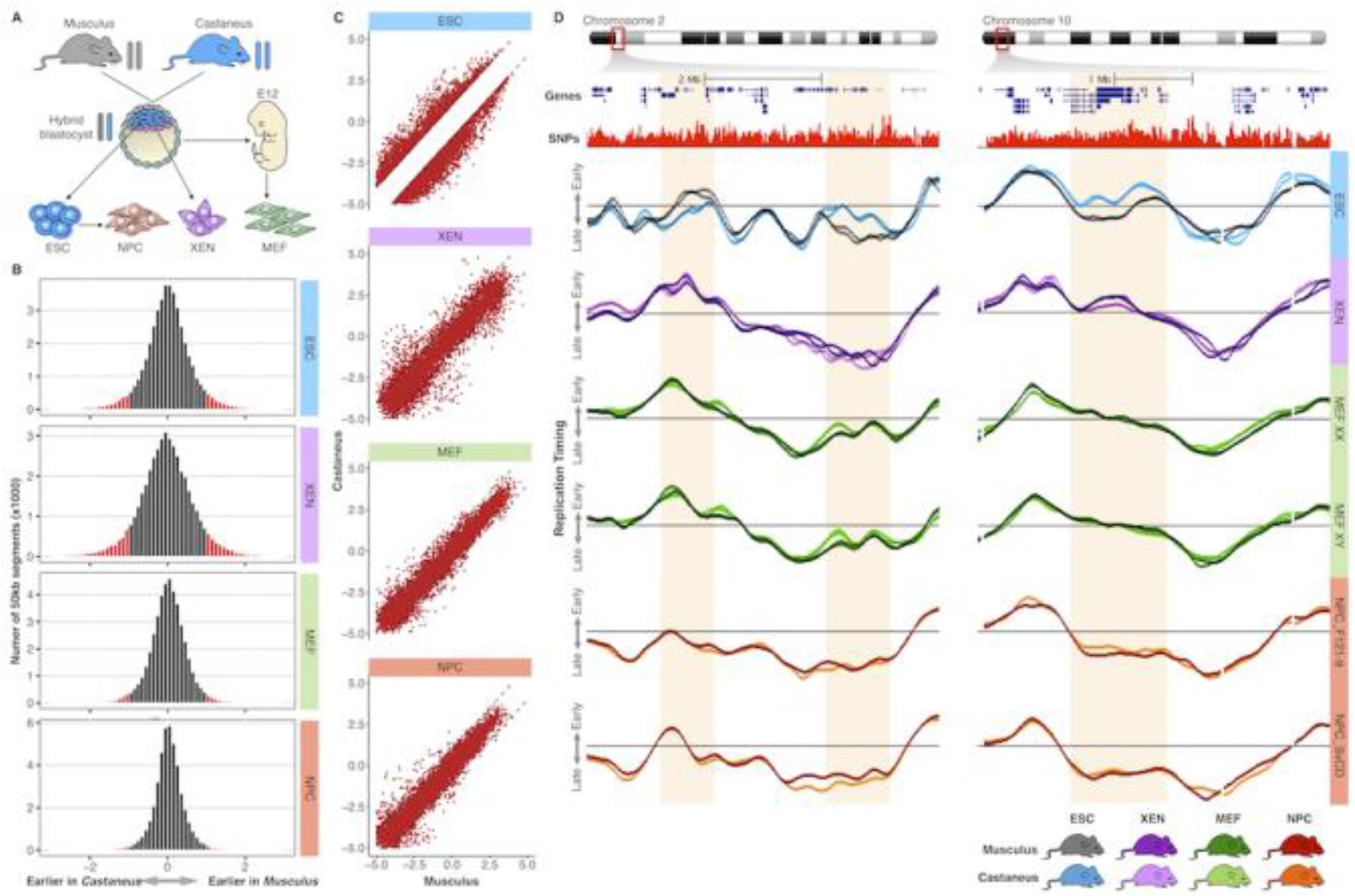
RT asynchrony is lost during differentiation (click for full resolution) (A) Distinct cell types were derived from hybrid crosses of *M. musculus musculus* x *M. musculus castaneus*. ESC and extraembryonic endoderm stem (XEN) cells were derived from hybrid blastocysts, mouse embryonic fibroblasts (MEF) from hybrid mouse embryos (E12-14) and neural precursor (NPC) cells were differentiated *in vitro* from hybrid ESC lines. (B) Histograms of RT differences between musculus (129/sv) and castaneus (CAST/Ei) genomes. Labeled in red are the changes higher than the differences between replicates of the same genome. (C) RT asynchronous regions in ESCs became synchronous in differentiated cell types. (D) Two exemplary chromosome regions display the loss of RT asynchrony in differentiated cell types. Two replicates of each sample are shown. Four distinct primary MEFs of distinct genders were obtained from different embryos, RT profiles of female MEFs (MEF XX) and male MEFs (MEF XY) are shown. Data from two independently differentiated NPCs derived from distinct ES cell lines (F121-9 and Cas129) are shown.

### Convergence of RT and 3D genome folding during differentiation

In order to understand the loss of RT asynchrony in differentiated cells we analyzed the changes in other genomic properties in NPCs including gene expression, chromatin accessibility and Hi-C compartments. Consistent with our findings in hybrid mouse ESCS, we found that early and late replicating domains correlate with Hi-C compartment A and B, respectively, and that early replicating regions have higher densities of chromatin interactions with shorter distances and are more accessible in comparison to late replicating regions (Supplementary Figure 10). Similarly, we found that RT correlates strongly with 3D chromatin organization genome-wide (Supplementary Figure 10). We then tracked the changes in chromatin interactions in the regions that replicate asynchronously in hybrid mESCs after differentiation to NPCs.

Consistent with the decrease in RT differences we found a convergence to similar chromatin organization with a decrease in differences between Hi-C compartments A and B (Supplementary Figure 10). In fact, we found that the chromosomal regions with allelic differences in RT and Hi-C compartments in stem cells, replicate synchronously and were organized within the same A/B compartment in NPCs (Figure 5).

**Figure 5.**
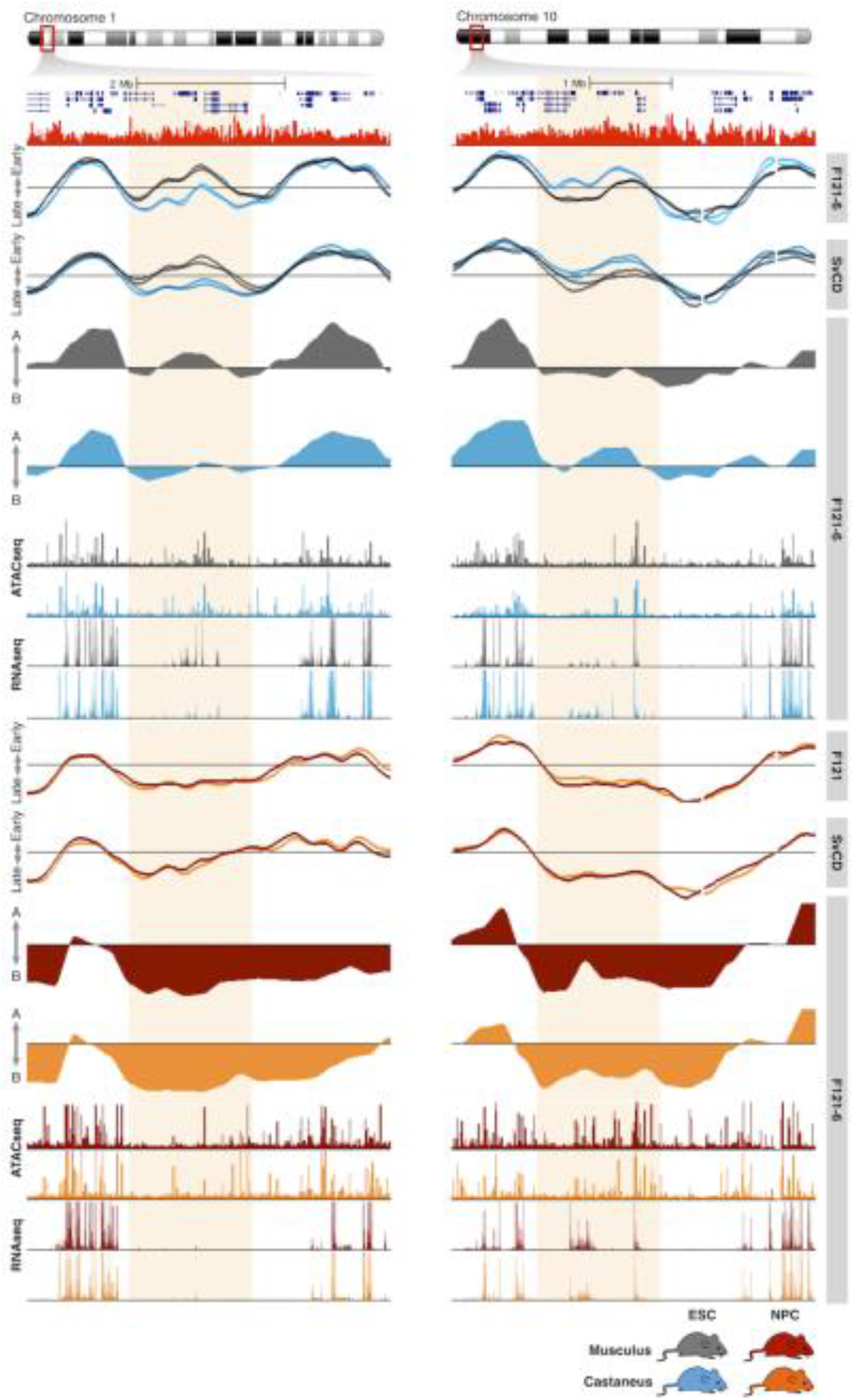
RT and genome organization control during development (click for full resolution) RT asynchrony loss is coordinated with a decrease in genome organization differences between alleles. Genome tracks of RT (two cell lines with opposite parental configuration), Hi-C eigenvectors, ATACseq and RNAseq are shown for mESCs (blue/grey) and for NPCs (red/orange). Differences in RT and Hi-C compartments are lost during differentiation to NPCs. Hi-C data was obtained from (Giorgetti et al. 2016).

## Discussion

In this study, we measured the allelic variation in genome organization and function exploiting the high SNP density between genomes of distinct mouse subspecies. Cell lines derived from crosses between *Mus musculus castaneus* and *Mus musculus musculus* subspecies exhibit a SNP on average every 150bp, rendering informative reads to determine the homologue chromosome of origin. To sort NGS reads into each homologue chromosome, we developed a new algorithm: *Haplotype-Assisted Read Parsing* (HARP), which can efficiently separate reads per genome based on the presence of SNPs. An advantage of HARP over other allele-parsing methods is the separated alignment of each genome that yields more reads as compared with alignment just to the reference genome. We measured allelic differences in RT, genome organization, gene expression and chromatin accessibility (Figure 1), providing a valuable resource of normal variation between homologues that can be leveraged as a reference framework for future studies. Normal variation between homologues would be particularly useful for genome editing studies to characterize gene function and chromatin architecture, where the generation of heterozygous mutants will provide an internal wild-type control for comparison and better assessment of the manipulation effects.

Previous studies found RT variation between homologues in 8-12% of the human genome (Mukhopadhyay et al. 2014; Bartholdy et al. 2015). Here, despite the 10 fold increase in SNP density, we found significant RT differences between mouse alleles in 12% of the autosomal genome in ES cells (Figure 2E). Interestingly, most of the genomic regions with allelic RT differences in mESCs overlap (67.4%) with developmentally regulated replication domains (Supplementary Figure 3), suggesting that RT asynchrony occurs at loci that change RT dynamically during development. Moreover, the overlap could be much higher, as the developmentally regulated regions were defined using only cell types with available data and additional genomic regions might change RT during cell fate commitment towards unexplored cell types. Hence, the overlap between RT asynchronous regions and developmentally regulated RDs is an underestimate but we cannot rule out that RT asynchrony can occur in regions that do not change RT during development. Consistent with previous studies (Ryba et al. 2010; Pope et al. 2014; Rivera-Mulia and Gilbert 2016b), we found that early replicating regions in both alleles correlated with Hi-C compartment A, higher gene expression and accessible chromatin when compared to late replicating regions (Figure 3). However, RT asynchrony correlated strongly with the 3D chromatin structure but no other genomic property (Figure 3).

In contrast with previous studies, where RT asynchrony was found associated with gene imprinting (Mukhopadhyay et al. 2014; Bartholdy et al. 2015), we found that allelic differences in RT did not correlate with gene imprinting (Supplementary Figure 5). Part of this could be quantitative; here we have imposed a stringent cut-off for only those regions that differ in RT more than the noise seen between replicate experiments. In addition, a caveat of previous studies was the lack of comparison of individuals with opposite parental configurations to confirm whether the allelic differences were linked to gene imprinting. Here, we took advantage of hybrid cell lines with opposite parental configurations and confirmed that the allelic differences in RT were not associated with parental imprinting (Supplementary Figure 5). We cannot rule out small differences in RT that are within the range of noise that may be detected by singlet-doublet FISH analysis (Selig et al. 1992; Boggs and Chinault 1997; Singh et al. 2003; Dutta et al. 2009). Our study does not, however, address random monoallelic asynchrony, which would require or single cell analyses. It is unlikely that random allelic asynchrony is occuring in mESCs, however, as several of our mESC lines are clonally derived.

Recent studies report gene expression changes and impaired differentiation capacity linked to culture conditions (MEFs in serum+LIF Vs. feeder-free in 2i+LIF) and associated with altered patterns of DNA methylation (Yagi et al. 2017; Choi et al. 2017). However, we found that RT asynchrony in mESCs is stable in all media conditions (Figure 2F and Supplementary Figure 4), and that hybrid mESCs maintained in any media conditions can be efficiently differentiated into neural precursors cells (NPCs).

X chromosome inactivation (XCI) in female cells occurs post-implantation during development in a random fashion in embryonic cell types; in extraembryonic tissues the paternal X is imprinted and inactivated (Takagi and Sasaki 1975; Escamilla-Del-Arenal et al. 2011). Consistently, early replication peaks were detected for both X chromosomes in mESCs but restricted to the musculus maternal allele in XEN cells (Supplementary Figure 11). Although, XCI occurs randomly in other cell types, we have shown previously that XCI can be detected as a decrease in RT values in differentiated cell types as compared to male cell lines (Hiratani et al. 2010). Consistently, we found a delay in RT for X chromosomes of all female differentiated cell lines (Supplementary Figure 11). Interestingly, we found that the musculus allele replicated later in all NPC and MEF cell lines, suggesting a sub-species skew in XCI (Supplementary Figure 11).

Finally, since almost all of the RT asynchrony has been reported in fully differentiated cell types, we also evaluated the allelic RT differences in XEN cells, primary MEFs and NPCs. Surprisingly, we found that the RT asynchrony observed in mESCs is lost during cell fate commitment (Figure 4), with more than 85 % of the loci showing allelic differences in RT in mESCs replicating synchronously in all differentiated cells. Moreover, although we found a few regions replicating asynchronously in differentiated cell types, allelic differences in RT were not associated with gene imprinting even in these differentiated cell types (Supplementary Figure 9). As discussed above, one possible explanation is that differences in RT associated with gene imprinting are lower than the significant differences considered in this study and additional replicates and/or single cell analysis would be required to identify the degree of RT variation of imprinted genes. Similarly, allelic differences in RT have been observed at randomly mono-allelically expressed genes (Donley et al. 2013). However, single cell sub-cloning of differentiated cell types will be required to assess the RT differences at mono-allelically expressed loci.

## Methods

### Cell culture

Hybrid mouse ES cell lines V6.5, F121, F121-6, F121-9, F123 and Cas129 (Figure 1b), were grown in two different conditions: a) on mitomycin-C-inactivated MEFs in ES media and b) feeder-free in 2i medium (see the Supplementary Table 1 for media composition details). For feeder free/serum free conditions cells were grown on 0.1% gelatin-coated dish in N2B27 media supplemented with 1 μM MEK inhibitor, 4.25 μM GSK3 inhibitor (1-Azakenpaullone), 2 mM glutamine and 1,000U/ml LIF (Cell Guidance Systems GFM200), 0.15 mM monothioglycerol. Hybrid XEN cells were derived and maintained as previously described {Kunath, et al 2005} in RPMI supplemented with 15% FBS (Life Tech), 100mM sodium pyruvate, 10mM B-mercaptoehanol and 200mM L-glutamine. Hybrid MEFs were grown in DMEM supplemented with 10% FBS, 1X NAAs, 2mM L-glutamine and 10mM B–mercaptoethanol. Neural precursor cells were derived from hybrid mouse ES cell lines with opposite parental configurations (F121-9 and Cas129. Hybrid mouse ESCs were grown in a monolayer culture at high density (1.5^105 cells/cm2) onto 0.1% (v/v) gelatin-coated 6-wells in serum-free medium ESGRO Complete Clonal Grade medium (CCGM). After 24 hours, cells were dissociated with 0.1% trypsin and then were plated onto 0.1% (v/v) gelatin-coated 10cm dishes at 1^104 cells/cm2 in RHB-A media (Takara), the media was changed every other day and NPCs were analyzed at day 11.

### RT libraries

Genome-wide RT profiles were constructed as previously described (Ryba et al. 2011; Marchal et al. 2017). Briefly, cells were pulse labeled with BrdU and separated into early and late S-phase fractions by flow cytometry, followed by DNA immunoprecipitation with anti-BrdU antibody. Sequencing libraries of BrdU-substituted DNA from early and late fractions were prepared by NEBNext Ultra DNA Library Prep Kit for Illumina (E7370). Sequencing was performed on Illumina-HiSeq 2500 by 100bp single end. Approximately 35 million reads per sample were generated.

### ATAC-Seq libraries

ATAC-Seq (Assay for Transposase-Accessible Chromatin using Sequencing) was performed as previously described (Buenrostro et al. 2013), starting with ~200,000 cells. Lysis was performed for 15 min on ice, and nuclei were collected by spinning at 600 g for 10 min at 4°C. Transposase from the Nextera DNA Sample Preparation Kit (Illumina) was added, and nuclei were incubated at 37°C for 30 min. Following DNA purification on a MinElute column (Qiagen), libraries were amplified using PCR reagents from the Nextera DNA Sample Preparation Kit and index primers from the Nextera Index Kit (Illumina). PCR amplification was performed using the following conditions: 72°C for 5 min; 98°C for 30 s; 10 cycles of 98°C for 10 s, 63°C for 30 s and 72°C for 3 min; and a final extension at 72°C of 5 min. Libraries were purified twice with PCR clean-up kit columns (Qiagen). Library concentration and size distribution were measured using the KAPA Library Quantification Kit (KAPA Biosystems) and the Agilent Bioanalyzer 2100. Libraries were sequenced (100 bp paired-end) on the HiSeq2500 platform (Illumina).

### Promoter Capture Hi-C libraries

Promoter Capture Hi-C was performed as previously described (Schoenfelder et al. 2015a), but using an in-nucleus ligation method for Hi-C library preparation (Nagano et al. 2015). Briefly, 3–4 × 10^7 cells were fixed in 2% formaldehyde for 10 min. Chromatin was digested with HindIII (NEB) overnight at 37°C, and Klenow (NEB) was used to label restriction sites with biotin-14-dATP (Life Technologies). In-nucleus ligation was performed by adding 25 units of T4 DNA ligase (Invitrogen) in T4 ligase buffer (NEB), supplemented with 100 μg/ml BSA, per 5 million cells starting material. The ligation reaction was incubated at 16°C overnight, and crosslinks were reversed by adding Proteinase K and incubating at 65°C overnight. Samples were treated with RNase A, and ligation products were purified by phenol/chloroform extraction. Biotin was removed from non-ligated Hi-C library fragment ends with T4 DNA polymerase (NEB), followed by phenol/chloroform extraction and sonication (Covaris E220). Hi-C library DNA was end-repaired with Klenow, T4 DNA polymerase, and T4 polynucleotide kinase (NEB). Library DNA was size selected with AMPure XP beads (Beckman Coulter) and 3’ ends were adenylated using Klenow exo–(NEB). Biotin-marked ligation products were bound to MyOne Streptavidin C1 Dynabeads (Life Technologies), and PE adapters (Illumina) were ligated. Bead-bound Hi-C DNA was amplified with 7 or 8 PCR amplification cycles using the PE PCR 1.0 and PE PCR 2.0 primers (Illumina), and amplified libraries were purified using AMPure XP beads (Beckman Coulter).

For the capture step, 500 ng of Hi-C library DNA were denatured by heating to 95°C for 5 min, and incubated at 65°C for 24 hrs with a custom-designed biotinylated RNA bait system targeting the ends of HindIII restriction fragments at 22,225 mouse gene promoters (Schoenfelder et al. 2015a) (Agilent Technologies). The Sure Select Target enrichment protocol (Agilent Technologies) was followed for biotin pull-down and washes, using MyOne Streptavidin T1 Dynabeads (Life Technologies). Captured Hi-C DNA was amplified with 4 PCR amplification cycles using the PE PCR 1.0 and PE PCR 2.0 primers (Illumina), and amplified libraries were purified using AMPure XP beads (Beckman Coulter). Promoter Capture Hi-C libraries were sequenced (100 bp paired-end) on the HiSeq1000 platform (Illumina).

### Nuclear RNAseq data

Previously reported strand-specific nuclear RNA-Seq libraries (Schoenfelder et al. 2015b) were re-sequenced (100 bp paired-end) on the HiSeq2500 platform (Illumina) and reads were merged with the published data for downstream analysis.

### Reads alignment and allele-specific assignment

Reads of quality scores above 30 were independently mapped to 129/Sv and *castaneus* reference genomes using bowtie2 and assigned to each allele using *HARP* (haplotype-assisted read parsing), a new computational algorithm to filter reads based on the presence of genome-specific SNPs (full documentation can be found in https://github.com/dvera/harp).

### Clustering analysis

Significant RT variable regions were defined as those regions with differences ≥ 1 in pairwise comparisons between all samples analyzed. Unsupervised hierarchical and k-means clustering analysis were performed using Cluster 3.0 (de Hoon et al. 2004) using un-centered correlation metrics and average linkage. Heatmaps and dendrograms were generated in JavaTreeView (Saldanha 2004).

## Data Access

### Datasets Availability

All datasets for replication timing, nuclear RNA-seq, ATAC-seq and PC-Hi-C generated in this study are deposited in the NCBI Gene Expression Omnibus database (GEO; http://www.ncbi.nlm.nih.gov/geo/), in the 4D Nucleome DCIC portal and in our laboratory database of (http://www.replicationdomain.org).

### Code Availability

Repli-Chip and Repli-Seq analysis was performed in R. RT datasets were normalized using the limma package in R and re-scaled to equivalent ranges by quantile normalization. Correlation analysis was performed using the corrplot package in R. Detailed computational pipeline for genome-wide measurement of RT has been published elsewhere (Ryba et al. 2011; Marchal et al. 2017). For allele-specific RT measurement we developed HARP (haplotype-assisted read parsing), a new computational algorithm (documentation in https://github.com/dvera/harp).

## Acknowledgments

We thank Ruth A. Didier for assistance with flow cytometry, Ferhat Ay for data processing advice.

This work has been supported by NIH grants GM083337 and DK107965 to DMG.

## Author’s contributions

JCRM and DMG conceived and designed the project. CD and JG generated the F1 hybrid cell lines. AD and PF generated PC-Hi-C, RNAseq and ATACseq datasets. JCRM, TS and CTG performed cell cultures and NPC differentiation. JCRM, TS and CTG collected the RT datasets; JCRM, DV and JZ performed data analysis and interpretation; JCRM and DMG wrote the manuscript.

## Supplemental Material

**Suppl. Table 1.** ES media composition

| Cell line | V6.5 | F121 | F121-6 | F121-9 | Cas129 | F123 |
| --- | --- | --- | --- | --- | --- | --- |
| DMEM | 200 mL | 500 mL | 500 mL | - | 500 mL | 500 mL |
| GMEM | - | - | - | 500 mL | - | - |
| FBS | 50 mL | 50 mL | 75 mL | 50 mL | 75 mL | - |
| KOSR | - | - | - | - | - | 50 mL |
| 100x NEAA | 3 mL | 5 mL | 5 mL | 5 mL | 5 mL | 6 mL |
| 100x Pyruvate | - | - | - | 5 mL | - | - |
| 100x Glutamine (200 mM) | 3 mL | 5 mL | 5 mL | 5 mL | - | - |
| 100x nucleosides* | 3 mL | - | - | - | - | - |
| 100x Glutamax | - | - | - | - | - | 6 mL |
| BME (FINAL) | 1 mM | 2.85 mM | 5 mM | 0.1 mM | 2.85 mM | 0.1 mM |
| 100x Pen/Strept | 3 mL | 5 mL | 5 mL | - | 5 mL | 6 mL |
| LIF | 10<sup>3</sup> U/mL | 10<sup>3</sup> U/mL | 10<sup>3</sup> U/mL | 10<sup>3</sup> U/mL | 10<sup>3</sup> U/mL | 10<sup>3</sup> U/mL |
| H2O | 37.5 mL | - | - | - | - | - |

**Suppl. Figure 1.**
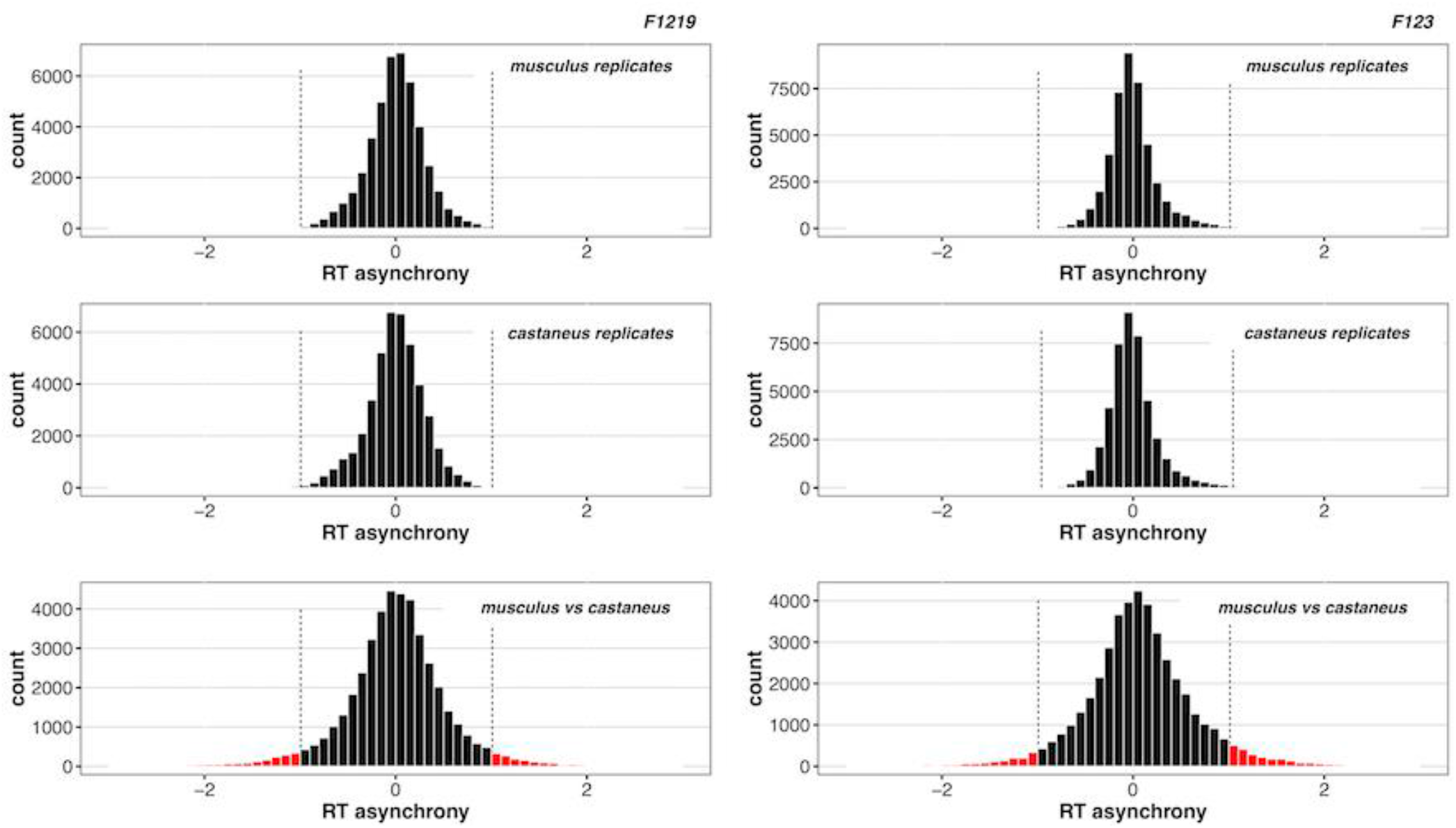
Replication timing asynchrony in hybrid mouse ESCs *castaneus* x *musculus* (<u>click for full resolution</u>). RT asynchrony in castaneus x musculus mES cell lines. Histogram of RT differences between replicates of alleles of the same genome and comparison between distinct genomes are shown.

**Suppl. figure 2.**
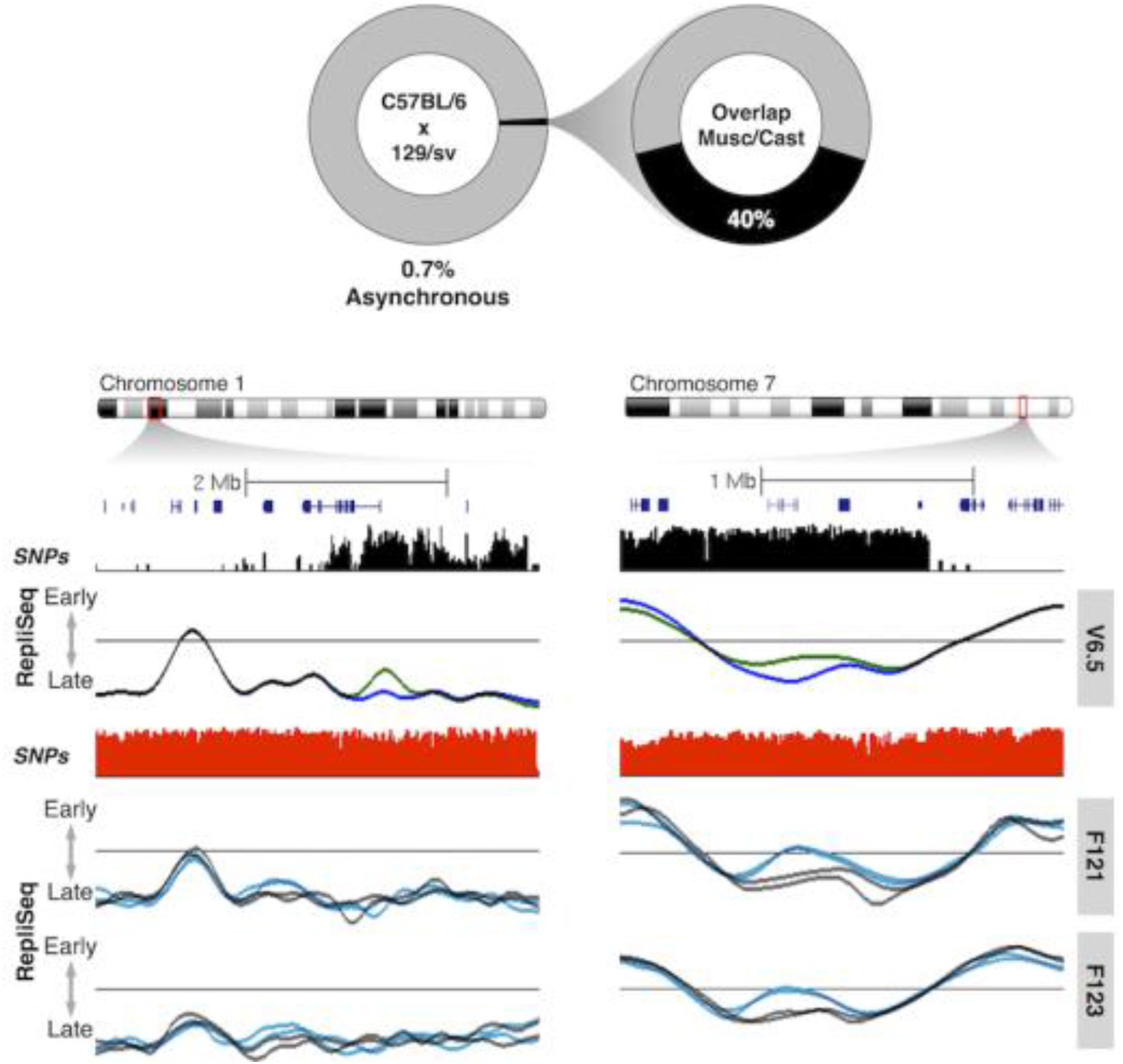
Replication timing asynchrony in hybrid mouse ESCs derived from inbred strains crosses (<u>click for full resolution</u>). RT asynchrony in V6.5 cell line (cross between C57BL/6 and 129/sv strains) and the overlap with the RT asynchrony observed in *musculus* X *castaneus* crosses. Exemplary regions are shown at the bottom. RT profiles show exemplary asynchronous region that are specific for the V6.5 (left) or that overlap with differences *musculus* x *castaneus* crosses (right).

**Suppl. figure 3.**
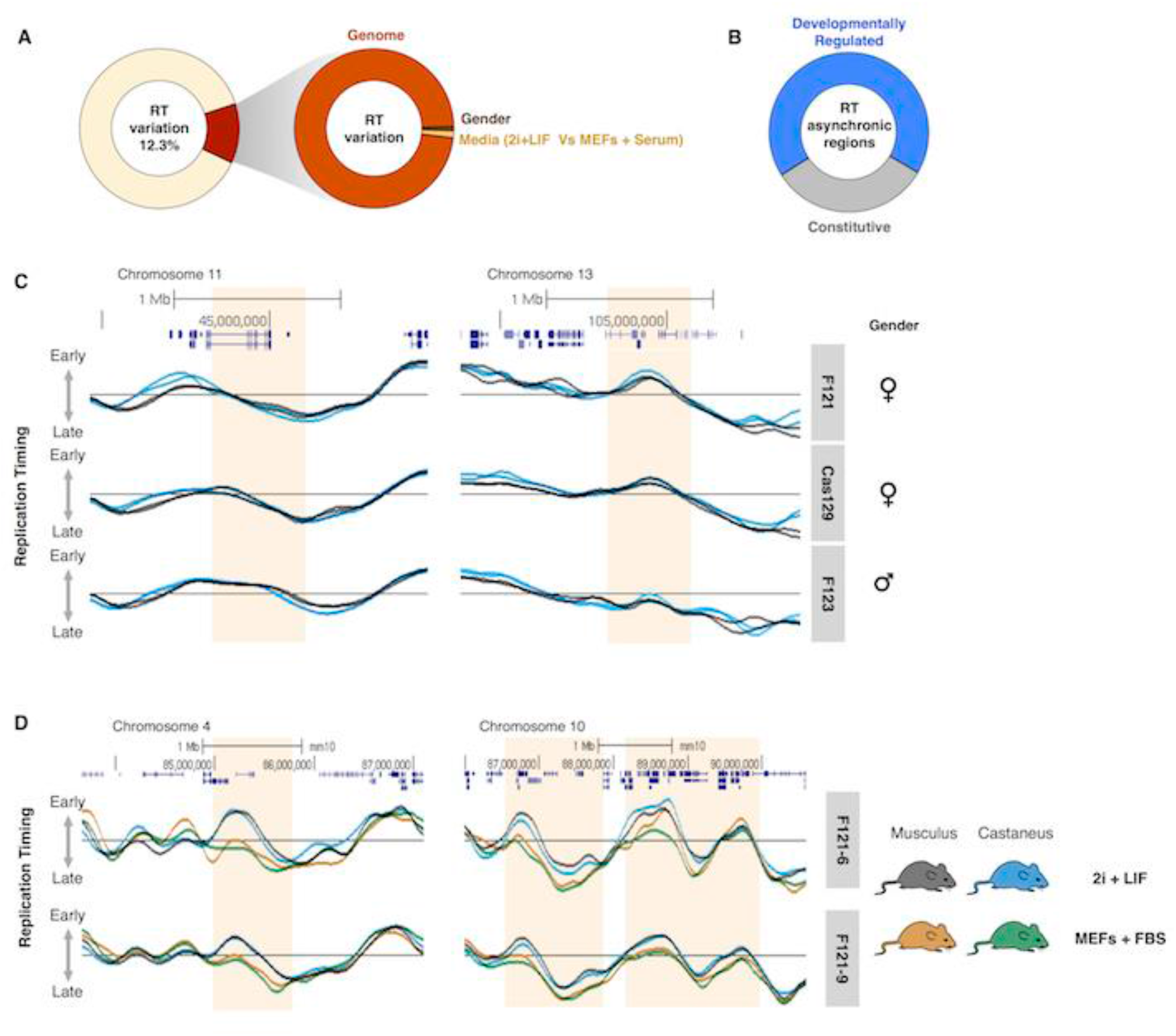
Replication timing variation is linked to sub-specie genomes (<u>click for full resolution</u>). (A) All RT variation across all cell lines was identified and sources quantified. 12% of the RT variation is associated to differences between sub-species genomes, 0.10% to gender differences and 0.17% to media conditions. No differences associated to parental configuration were found. (B) Asynchronously replicating domain in mESCs are enriched in genomic regions that are change RT during development. (C) Exemplary regions with differences between genders. Left RT profiles show a region that replicates earlier in male cell lines, while right RT profiles show a region that replicate earlier in females. (D) Exemplary regions showing RT differences linked to media conditions.

**Suppl. figure 4.**
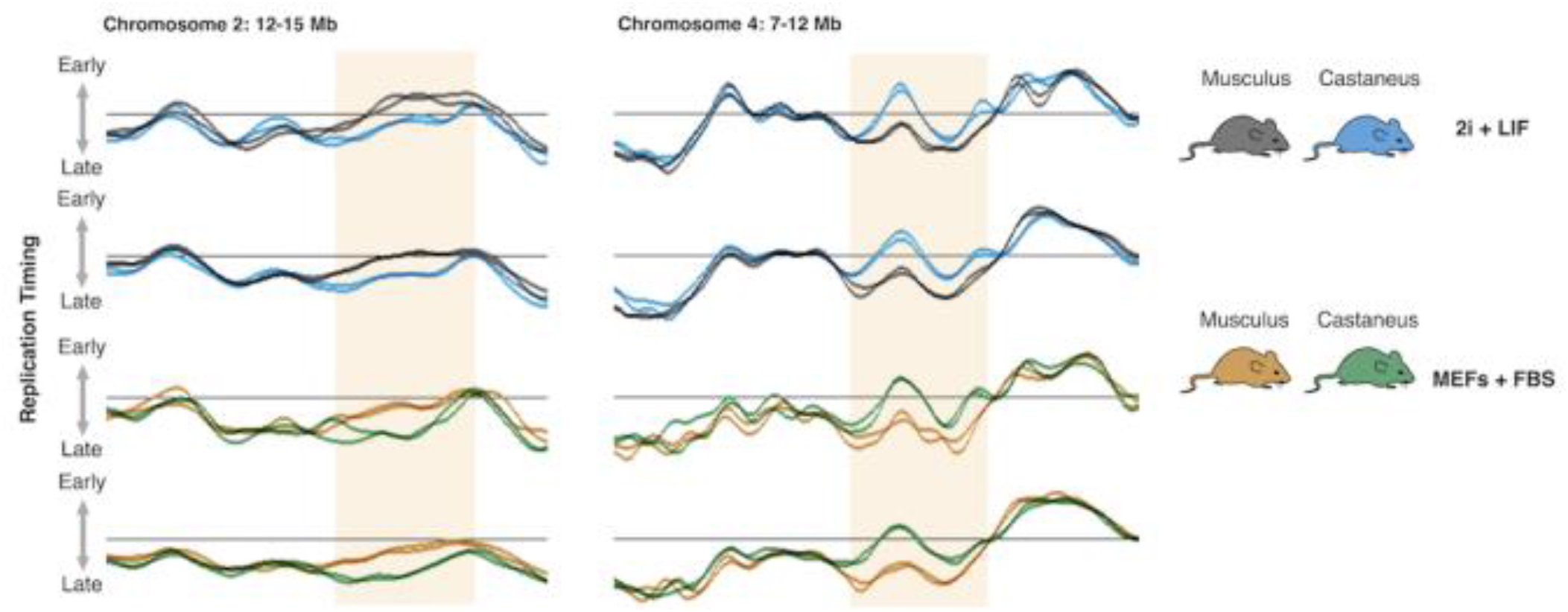
Replication timing asynchrony is maintained in different growing conditions (<u>click for full resolution</u>). Exemplary RT asynchronous regions from two different mES cell lines cultured in different media conditions.

**Suppl. figure 5.**
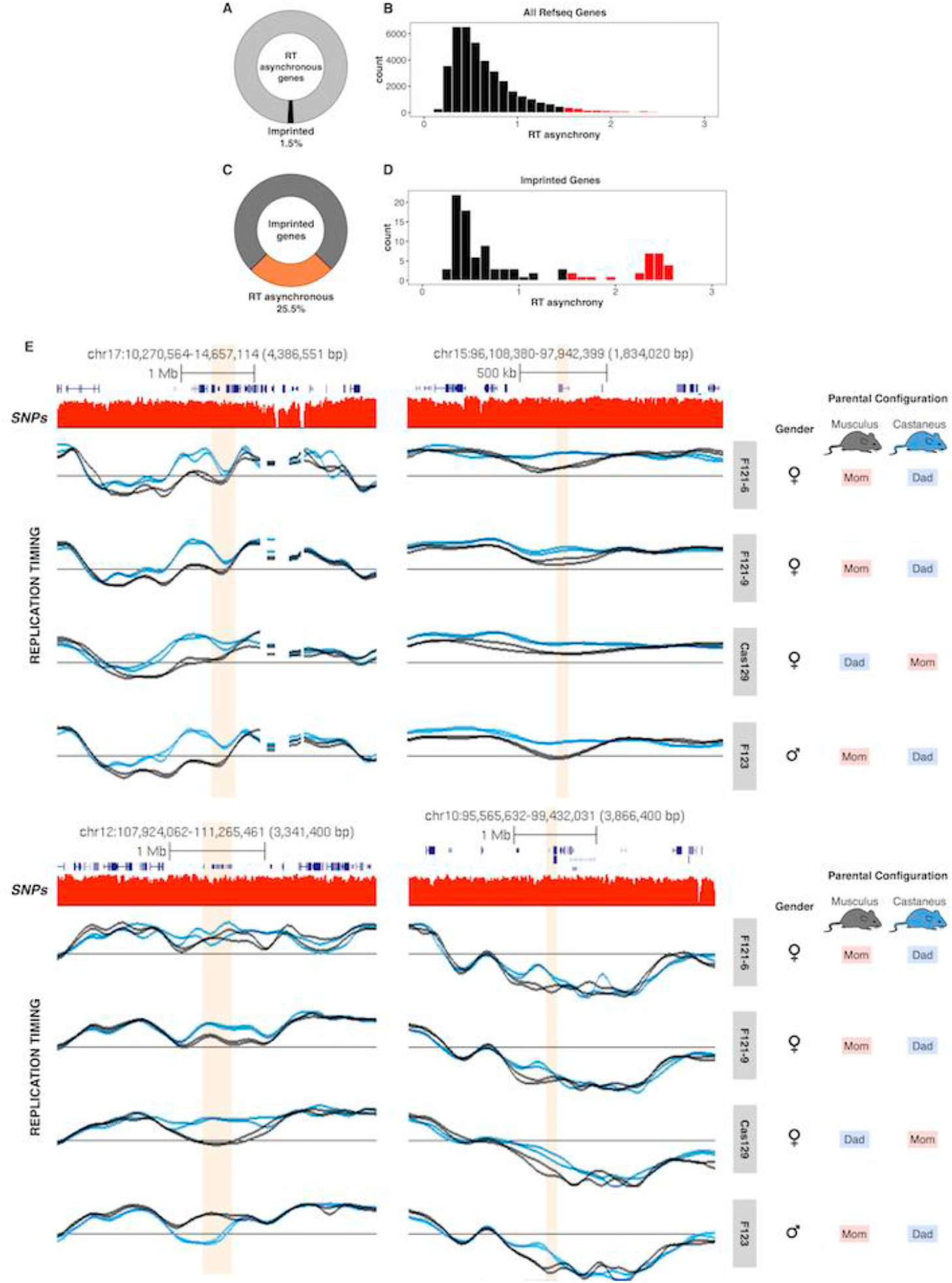
Replication timing asynchrony is not associated to gene imprinting (<u>click for full resolution</u>). (A) RT asynchrony is not enriched at imprinted genes in hybrid mouse ES cells. RT values at the transcription start sites (TSS) of all RefSeq genes were obtained and asynchronous genes identified. Only 1.5% of the RT asynchronous genes are imprinted. (B) RT asynchrony across all RefSeq genes. (C) Only a small fraction (25.5%) of the imprinted genes replicate asynchronously in hybrid mouse ES cells. (D) RT asynchrony in imprinted genes. (E) Exemplary RT profiles at 4 genomic regions that include the imprinted genes with the highest difference in RT. Imprinted genes are highlighted in each plot. Differences in RT are linked to the respective genomes rather than gene imprinting. List of imprinted genes was extracted from the imprinted genes database: www.geneimprint.com

**Suppl. figure 6.**
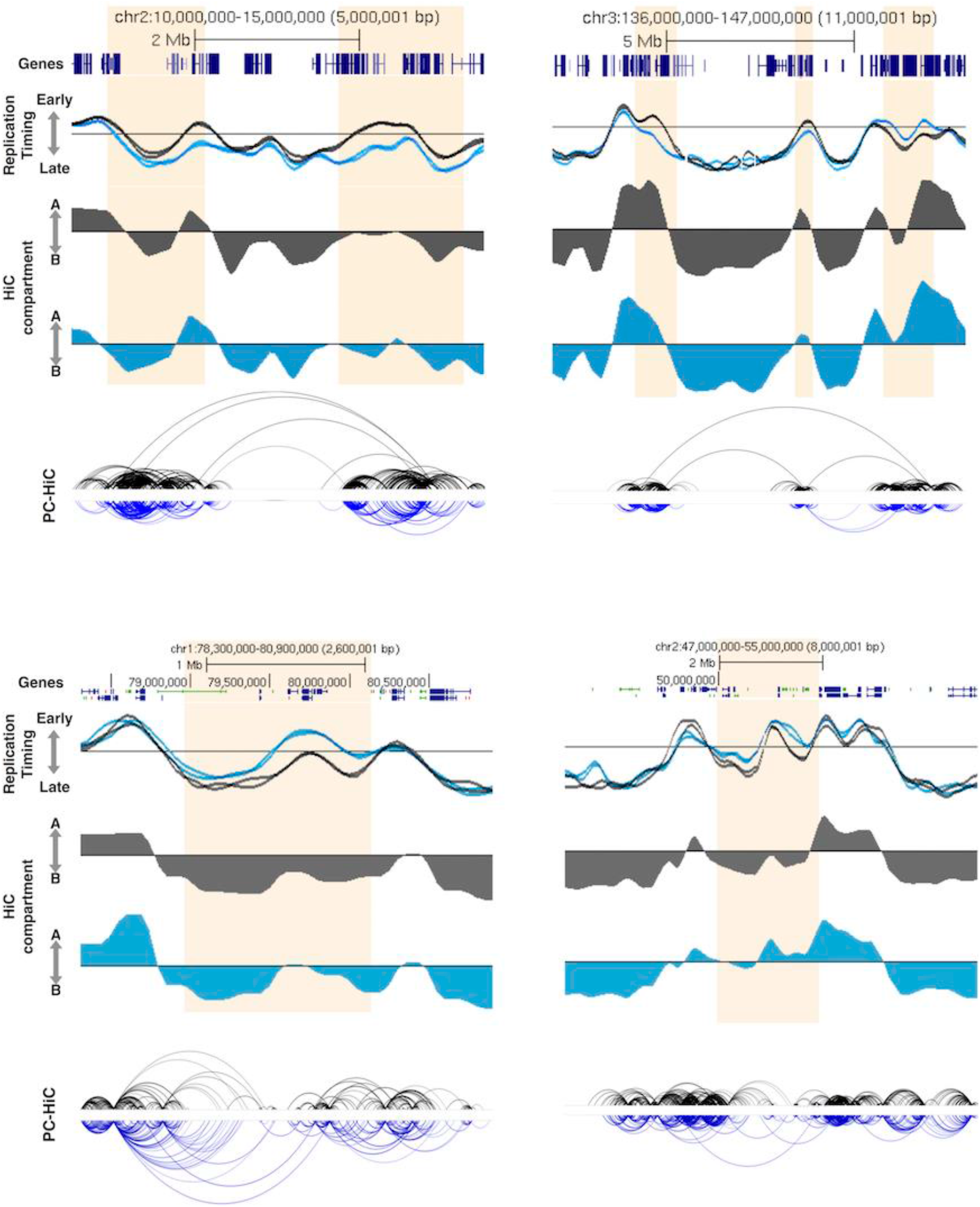
RT asynchrony is linked to long-range enhancer-promoter interactions (<u>click for full resolution</u>). Long-range enhancer interactions are restricted to the allele that replicates earlier. In the two exemplary regions at the top long-range interactions connecting the asynchronous domain with other distant early replication domains are restricted to the musculus allele (grey), while the regions at the bottom show long-range interactions restricted to the castaneus allele (blue).

**Suppl. figure 7.**
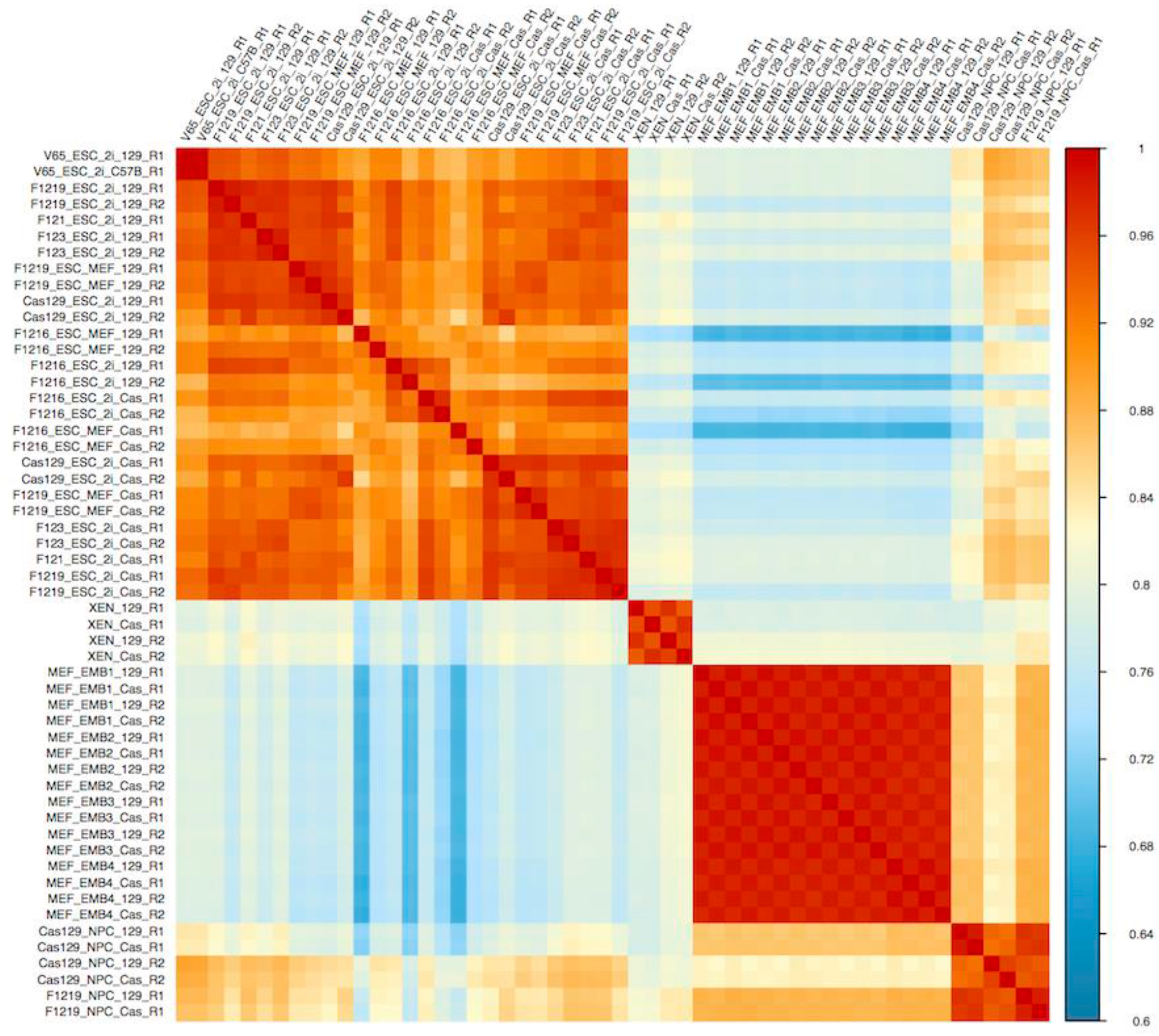
Correlation matrix of RT programs from all hybrid mouse cell types (<u>click for full resolution</u>). Genome-wide correlation of RT programs from all hybrid mouse cell types including ES cells, XEN cells, MEFs and NPCs confirms that RT is cell type specific. Moreover, consistent with loss of RT asynchrony upon cell fate commitment, stronger correlations between alleles, replicates and cell lines were observed in all differentiated cells as compared to ESCs.

**Suppl. figure 8.**
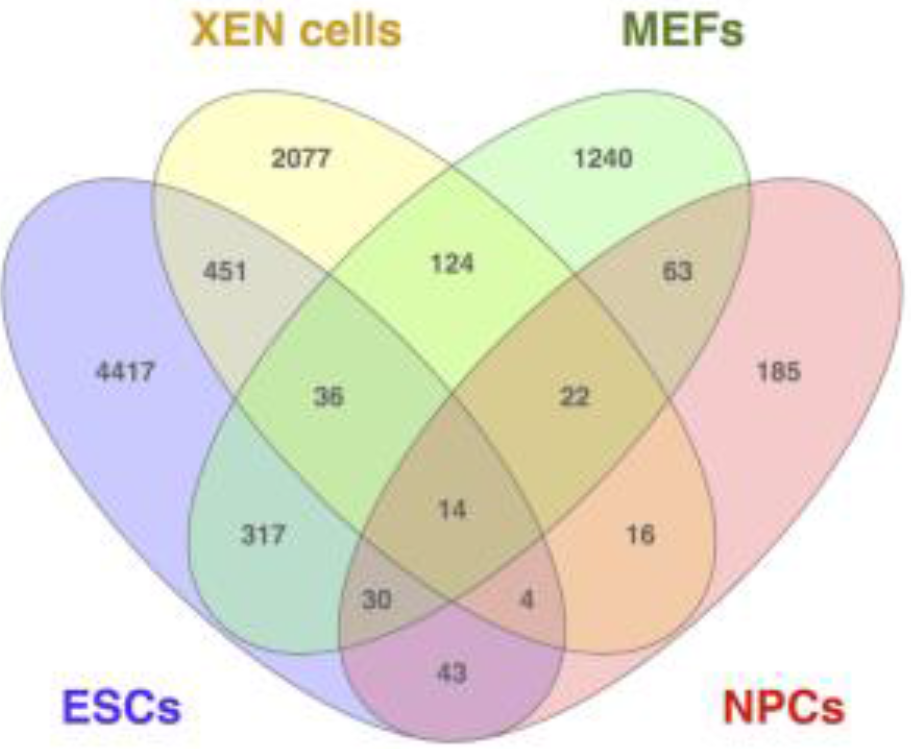
Distinct regions replicate asynchronously in each hybrid mouse cell type (<u>click for full resolution</u>). Overlap analysis of RT asynchronous regions identified in all hybrid mouse cell types.

**Suppl. figure 9.**
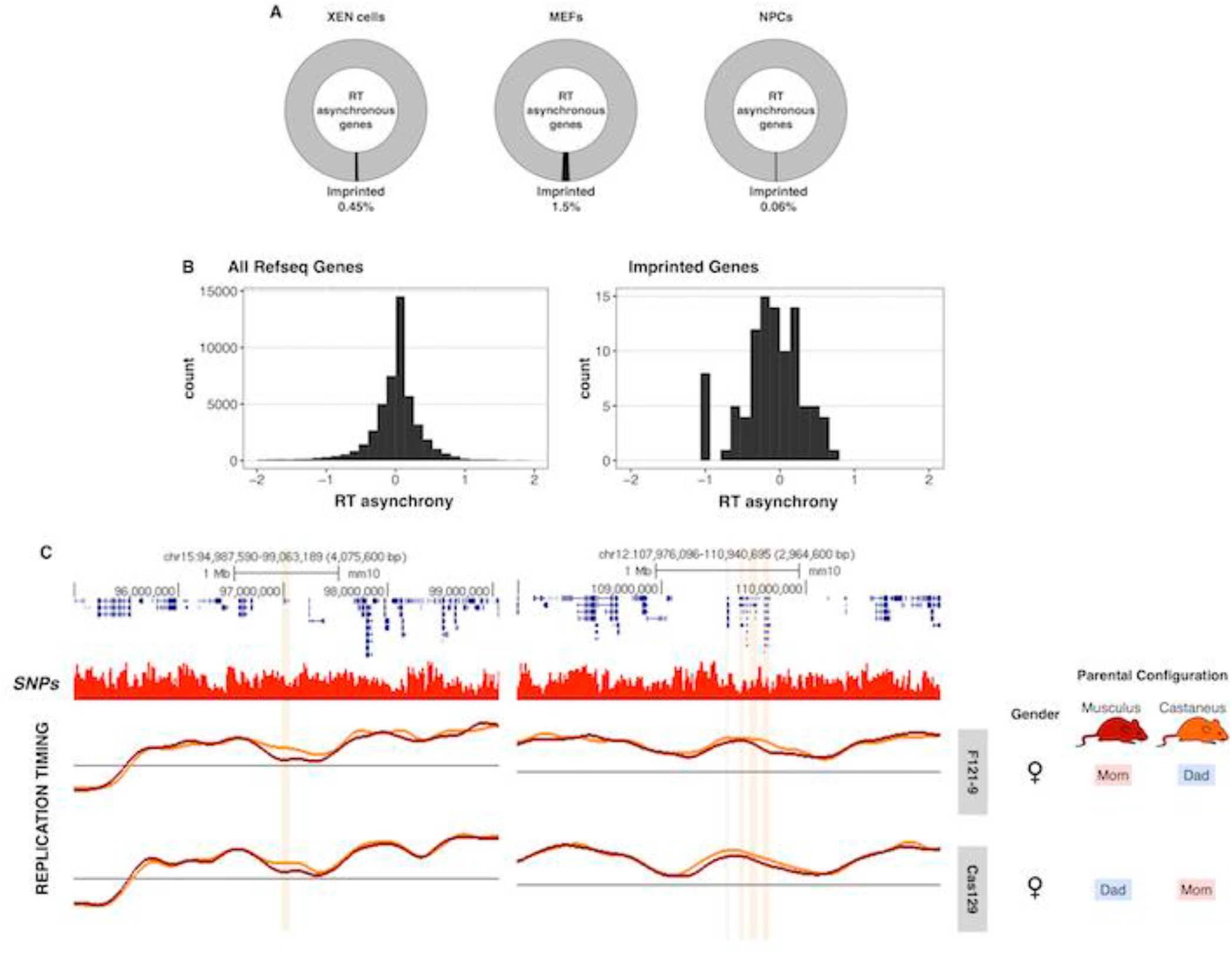
RT asynchrony is not associated with gene imprinting in differentiated cells (<u>click for full resolution</u>). (A) RT asynchronous genes and gene imprinting in differentiated cell types. RT values at the transcription start sites (TSS) of all RefSeq genes were obtained and asynchronous genes identified. RT asynchronous genes are not associated with gene imprinting. (B) RT asynchrony across all RefSeq genes and for imprinted genes in NPCs. (C) Exemplary RT profiles of the 2 imprinted loci with the highest RT differences in NPCs derived from hybrid mES cell lines with opposite parental configuration.

**Suppl. figure 10.**
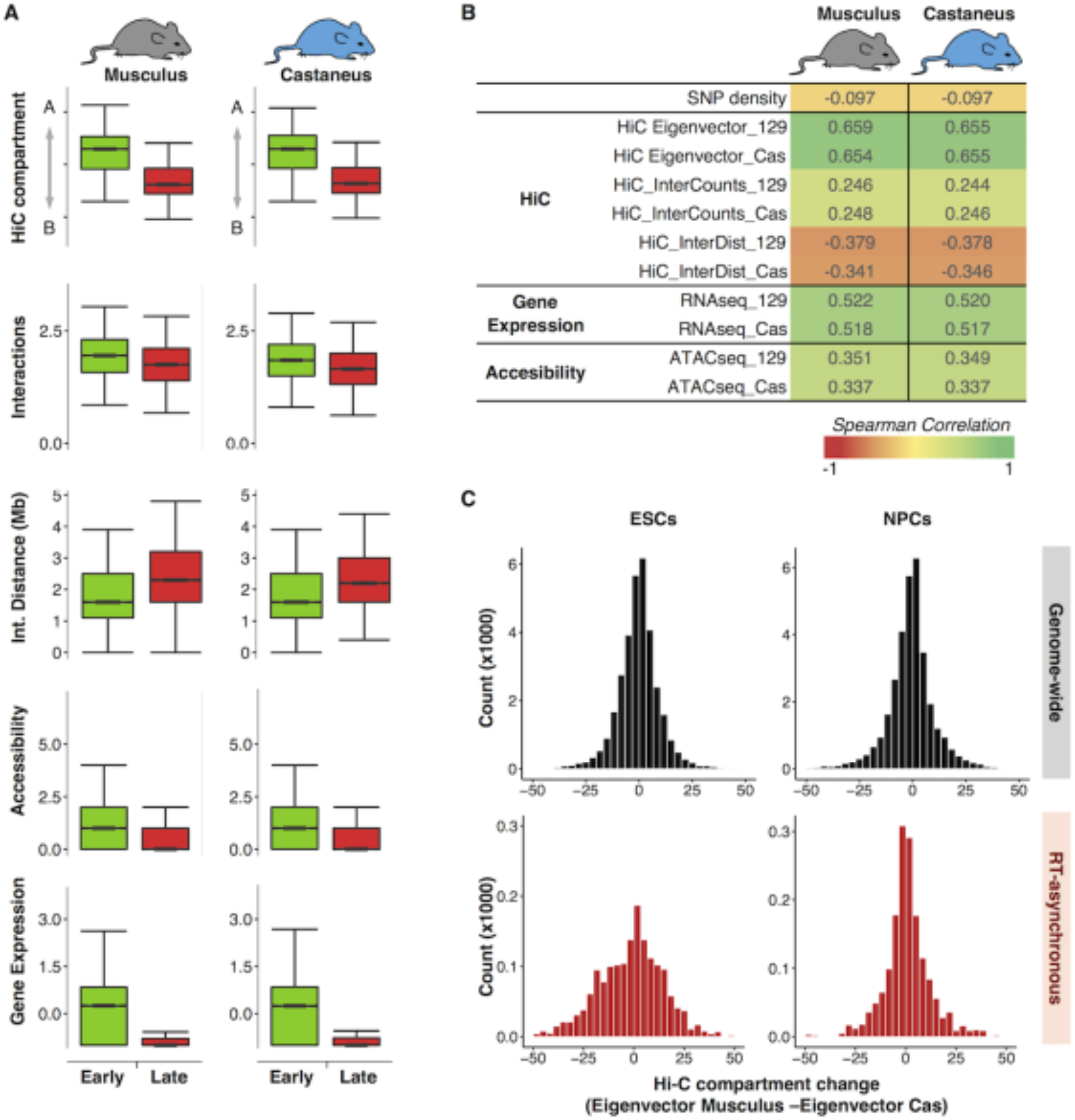
RT and genome organization in NPCs (<u>click for full resolution</u>). (A) Genome organization, chromatin accessibility and gene expression of RT synchronous regions that replicate either early or late during S-phase in NPCs. (B) Spearman correlation values of RT and distinct genomic features per genome. (C) Histograms of Hi-C eigenvector differences between genomes in hybrid ESCs and NPCs. Top panels show the differences genome-wide, bottom panels show the differences for the regions that replicate asynchronously in ESCs.

**Suppl. figure 11.**
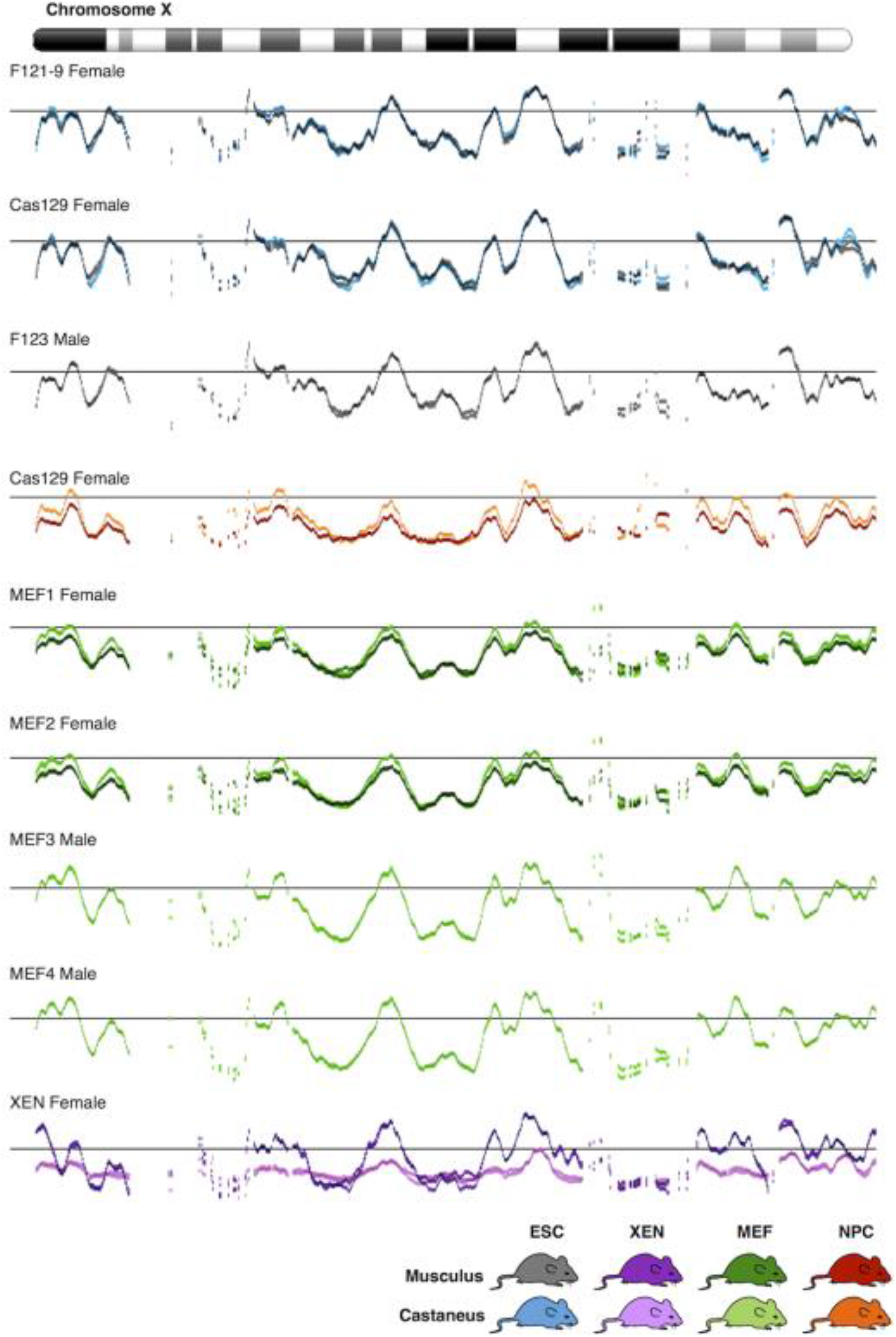
RT profiles of the chromosome X in distinct cell types derived from hybrid mouse crosses (<u>click for full resolution</u>).

